# An Interpretable Bayesian Clustering Approach with Feature Selection for Analyzing Spatially Resolved Transcriptomics Data

**DOI:** 10.1101/2023.05.10.540273

**Authors:** Huimin Li, Xi Jiang, Lei Guo, Yang Xie, Lin Xu, Qiwei Li

## Abstract

Recent breakthroughs in spatially resolved transcriptomics (SRT) technologies have enabled comprehensive molecular characterization at the spot or cellular level while preserving spatial information. Cells are the fundamental building blocks of tissues, organized into distinct yet connected components. Although many non-spatial and spatial clustering approaches have been used to partition the entire region into mutually exclusive spatial domains based on the SRT high-dimensional molecular profile, most require an ad-hoc selection of less interpretable dimensional-reduction techniques. To overcome this challenge, we propose a zero-inflated negative binomial mixture model to cluster spots or cells based on their molecular profiles. To increase interpretability, we employ a feature selection mechanism to provide a low-dimensional summary of the SRT molecular profile in terms of discriminating genes that shed light on the clustering result. We further incorporate the SRT geospatial profile *via* a Markov random field prior. We demonstrate how this joint modeling strategy improves clustering accuracy, compared with alternative state-of-the-art approaches, through simulation studies and two real data applications.

## 1. Introduction

Recent advances in spatially resolved transcriptomics (SRT) technologies have enabled the comprehensive molecular and spatial characterization of single cells. Understanding the spatial organization of cells, together with their molecular profiles (e.g., mRNA and protein abundances), provides valuable insight into their underlying biological functions, such as embryo development (Satija et al., 2015) and tumor progression (De Bruin et al., 2014). Spatial analysis holds enormous potential for deepening our understanding of biomedicine and has led to the increasingly common application of SRT technologies in various fields, such as cancer research and developmental biology (Marx, 2021), resulting in an explosive generation of SRT data. Current SRT techniques are either imaging-based by way of single-molecule fluorescence *in situ* hybridization (FISH), such as seqFISH (Eng et al., 2019; Lubeck et al., 2014) and STARmap (Wang et al., 2018), or sequencing-based by way of spatial barcoding, such as spatial transcriptomics (ST) (Ståhl et al., 2016) and Slide-seq (Rodriques et al., 2019). Imaging-based methods can measure hundreds of genes across a large number of individual cells, while sequencing-based methods can measure tens of thousands of genes at spatial locations, also known as spots, consisting of multiple cells. In summary, these SRT technologies integrate geospatial and molecular profiles to enable simultaneous measurement and spatial mapping of gene expression over a tissue slide, providing a comprehensive understanding of cellular organization and function.

The emergence of SRT technologies provides new opportunities to investigate research questions. Clustering is an essential step in the analysis of SRT data. It is crucial for downstream analyses, such as cell typing and differential expression analysis, providing insight into underlying biological processes. Non-spatial clustering methods such as *k*-means and the Louvain method (Blondel et al., 2008) may lead to scattered clusters, as the algorithm only takes gene expression as input and ignores spatial information. Recently, several methods have been developed to incorporate available spatial information to account for spatial correlation of gene expression: stLearn (Pham et al., 2020) uses deep learning features extracted from the histology image and the expression of spatially neighboring spots to account for spatial correlation in gene expression; BayesSpace (Zhao et al., 2021) uses a Bayesian approach by employing a Markov random field (MRF) prior, encouraging neighboring spots to belong to the same cluster; SpaGCN (Hu et al., 2021) employs a graph convolutional network approach that combines gene expression, spatial location, and histology to perform clustering; SC-MEB (Yang et al., 2022) conducts spatial clustering with a hidden MRF using an empirical Bayes approach; and DR-SC (Liu et al., 2022) simultaneously performs dimension reduction and clustering via a unified statistically-principled method.

Although these spatial clustering methods can perform clustering analysis on SRT data, they all have clear limitations. Firstly, all of these methods must transform counts in the SRT molecular profile to continuous levels for the sake of convenient statistical modeling. However, this extra step does not truly reflect the underlying data generation mechanism and may cause information loss (Sun et al., 2020). Secondly, SRT data commonly contain a substantial number of zeros (e.g., 60% - 90%), an issue that may significantly reduce statistical power and is handled by none of the aforementioned methods. And last but not least, to overcome the curse of dimensionality, most of them require linear or non-linear dimension reduction procedures to obtain a low-dimensional representation of data before conducting clustering. Such procedures include principal component analysis (PCA), t-distributed stochastic neighbor embedding (tSNE) (Van der Maaten and Hinton, 2008) and uniform manifold approximation and projection (UMAP) (McInnes et al., 2018). This introduces two shortcomings: 1) They require selecting less interpretable leading components compared with the original features, and 2) their results are sub-optimal due to information loss.

Motivated by Li et al. (2017), which proposed a zero-inflated Poisson-Dirichlet process mixture model with feature selection for text analysis, we developed a Bayesian finite mixture model with an MRF prior to simultaneously cluster spots or cells and identify the associated discriminating genes for SRT data analysis, named <u>Bayes</u>ian <u>c</u>lustering <u>a</u>pproach with <u>f</u>eature <u>s</u>election (BayesCafe). BayesCafe directly models the molecular profile of SRT data using a zero-inflated negative binomial (ZINB) mixture model to account for the zero-inflation and over-dispersion observed in SRT data and avoids choosing an *ad-hoc* data normalization method. To increase interpretability, BayesCafe utilizes a feature selection approach that offers low-dimensional representations of the SRT data in terms of a list of discriminating genes. This enables any gene-wise downstream analyses. Furthermore, BayesCafe employs an MRF prior to integrate the geospatial profile of SRT data to improve clustering accuracy. We demonstrated the advantages of BayesCafe through a comprehensive simulation study that considers various spatial patterns and zero-inflation settings. In addition, we applied BayesCafe to one sequencing-based and one imaging-based SRT dataset, showing improved clustering accuracy compared to existing methods.

The remainder of the paper is organized as follows: Section 2 introduces the zero-inflated hierarchical mixture model and discusses its parameter structure. In Section 3, we present the Markov chain Monte Carlo (MCMC) algorithms. Sections 4 and 5 evaluate BayesCafe on simulated and two real datasets. The final section concludes the article and discusses potential future research directions.

## 2. Model

In this section, we introduce BayesCafe for spatial clustering and feature selection. The overall workflow is depicted in Figure 1. Supplementary Figure S1 and Table S1 summarize the graphical and hierarchical formulation of the proposed model.

**Figure 1:**
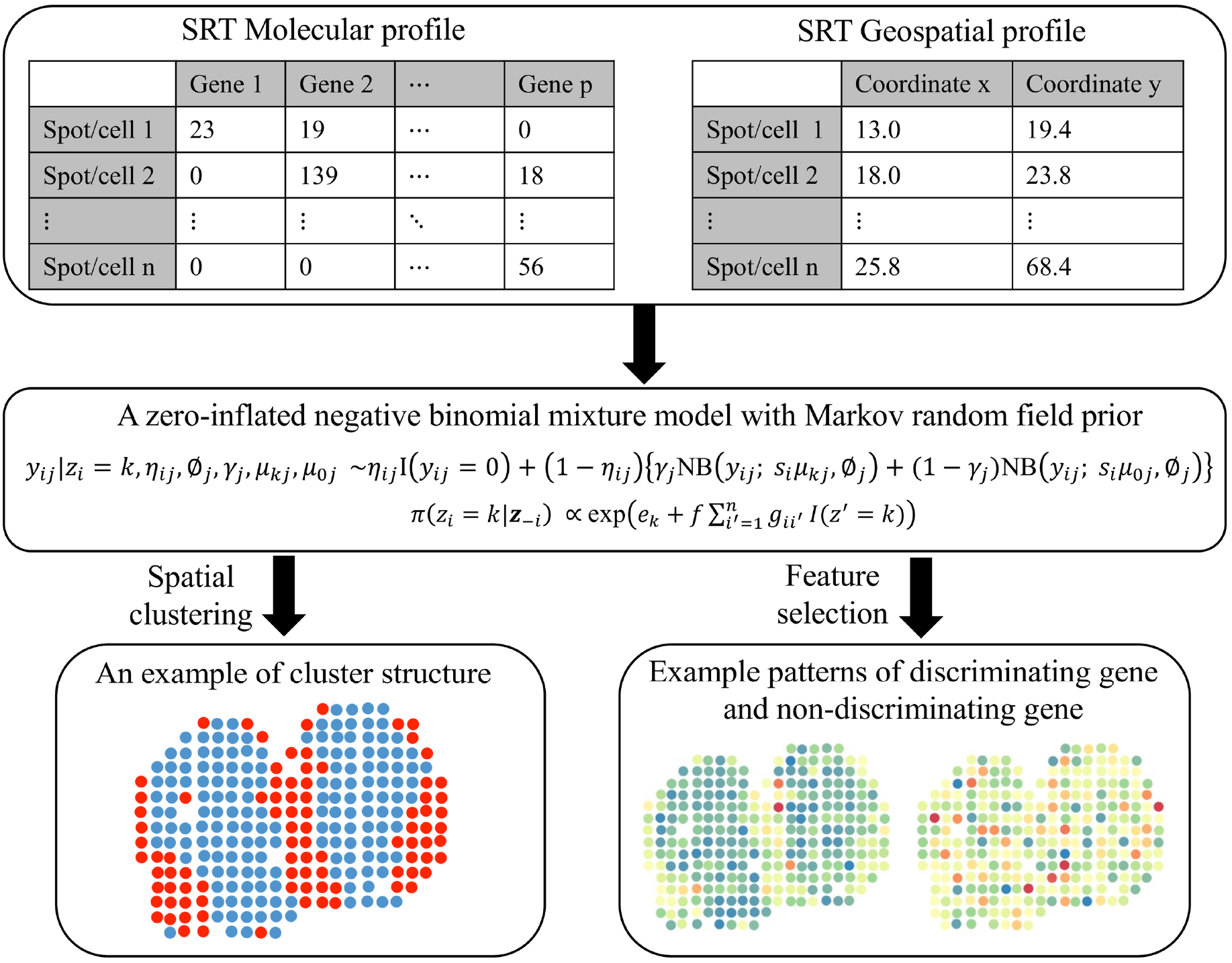
The schematic diagram of the proposed BayesCafe model.

We begin by summarizing observed data as follows: Let a *n*-by-*p* count matrix ***Y*** denote the molecular profile generated by an SRT technique. Each entry *y*_*ij*_ *∈* ℕ, *i* = 1, *…, n, j* = 1, *…, p*, is the read count observed in sample point *i* (i.e., spot or cell) for gene *j*. Let a *n*-by-2 matrix ***T*** represent the geospatial profile, where each row ***t***_*i·*_ = (*t*_*i*1_, *t*_*i*2_) *∈* ℝ^2^ gives the coordinates of sample point *i* in a compact subset of the two-dimensional Cartesian plane (see examples in Figure 1).

### 2.1 Modeling the SRT Molecular Profile via a ZINB model

Previous studies on scRNA-seq data analysis have suggested that accounting for the large proportion of zeros in the model can lead to a substantial improvement in model fitting and accuracy of identifying differentially-expressed genes (Finak et al., 2015; Lun et al., 2016); therefore, we start by considering a ZINB model to model the read counts:

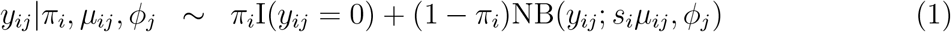

where we use I(*·*) to denote the indicator function and NB(*μ, ϕ*), *μ, ϕ >* 0 to denote a negative binomial (NB) distribution with expectation *μ* and dispersion 1*/ϕ*. With this NB parameterization, the probability mass function (p.m.f) is written as 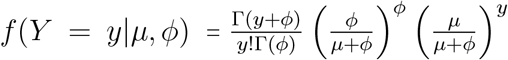 with variance Var(*Y*) = *μ* + *μ*^2^*/ϕ*, thus allowing for over-dispersion.

A small value of *ϕ* indicates a large variance-to-mean ratio, while a large value approaching infinity reduces the NB model to a Poisson model with the same mean and variance. In this model, we constrain one component of the mixture model to degenerate at zero, thereby allowing for zero-inflation. The sample-specific parameter *π*_*i*_ *∈* (0, 1) can be treated as the proportion of extra zero (i.e., false zero or structural zero) counts at sample point *i*.

The mean in the NB distribution is further decomposed into two multiplicative effects: the size factor *s*_*i*_ *∈* ℝ+ and the normalized expression level *μ*_*ij*_ *∈* ℝ+. The collection ***s*** = (*s*_1_, …, *s*_*n*_)^*⊤*^ reflects many nuisance effects across sample points, including but not limited to: 1) Reverse transcription efficiency; 2) Amplification and dilution efficiency; and 3) Sequencing depth. After adjusting for the global sample-specific effect, *μ*_*ij*_ can be regarded as the normalized expression level of gene *j* observed at sample point *i*. We follow (Sun et al., 2020) to set *s*_*i*_ proportional to the summation of the total number of read counts across all genes observed at sample point *i*, combined with a constraint of 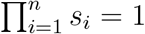 This results in 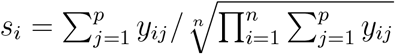 After accounting for zero-inflation (*via π*_*i*_’s), over-dispersion (*via ϕ*_*j*_’s), and sample point heterogeneity (*via s*_*i*_’s), our modeling approach produces a denoised version of gene expression levels, represented by *μ*_*ij*_’s.

To create an environment conducive to model fitting, we rewrite (1) by introducing a latent indicator variable *η*_*ij*_:

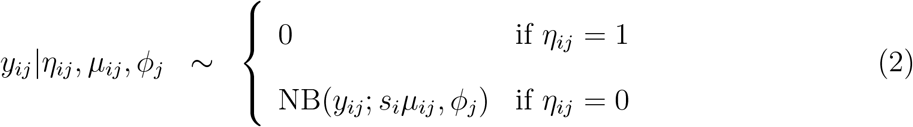

We impose an independent Bernoulli prior for *η*_*ij*_, i.e. *η*_*ij*_ ∼ Bern(*π*_*i*_), which can be further relaxed by formulating a Be(*a*_*π*_, *b*_*π*_) hyperprior on *η*_*ij*_, leading to a Beta-Bernoulli prior of *η*_*ij*_ with expectation *a*_*π*_*/*(*a*_*π*_ + *b*_*π*_). Setting *a*_*π*_ = *b*_*π*_ = 1 results in a weakly informative prior on *π*_*i*_. We assume a gamma prior for all dispersion parameter *ϕ*_*j*_’s, i.e. *ϕ*_*j*_ ∼ Ga(*a*_*ϕ*_, *b*_*ϕ*_) and suggest small values such as *a*_*ϕ*_ = *b*_*ϕ*_ = 0.001 for a weakly informative setting.

### 2.2 Clustering Spots/Cells While Identifying Discriminating Genes via a ZINB Mixture Model

When dealing with high-dimensional datasets, many features can provide very little information, and the inclusion of unnecessary features can complicate or even mask cluster recovery; therefore, we envision that only some features are relevant to discriminate *n* spots or cells into *K* distinct clusters. To identify these discriminating genes, we first introduce a latent binary vector for gene ***γ*** = (*γ*_1_, *…, γ*_*p*_)^*⊤*^, with *γ*_*j*_ = 1 if gene *j* is differentially expressed among *K* clusters (i.e., discriminating genes), and *γ*_*j*_ = 0 otherwise. Conditioning on *η*_*ij*_ = 0, we can write our model as follows:

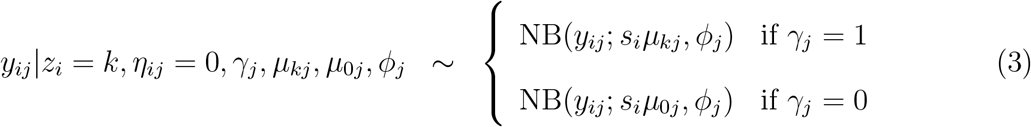

If *η*_*ij*_ = 1, we assume *y*_*ij*_ = 0 irrespective of *γ*_*j*_ with *i* = 1,… *n* and *j* = 1,… *p*. We utilize the common independent Bernoulli prior for *γ*_*j*_ with a hyperparameter *ω*, that is, *γ*_*j*_|*ω* ∼ Bern(*ω*), which is equivalent to a binomial prior on the number of discriminating genes, i.e.,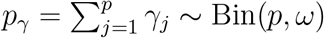 The hyperparameter *ω* can be elicited as the proportion of genes expected *a priori* to be in the discriminating gene set. This prior assumption can be further relaxed by formulating a beta hyperprior Be(*a*_*ω*_, *b*_*ω*_) on *ω*, which produces a Beta-Binomial prior on the number of discriminating genes *p*_*γ*_ with expectation *pa*_*ω*_*/*(*a*_*ω*_ + *b*_*ω*_). We set a vague prior on *ω* with a constraint of *a*_*ω*_ + *b*_*ω*_ = 2 (Tadesse et al., 2005).

For the task of cluster assignment, an auxiliary set of cluster allocation variables ***z*** = (*z*_1_, *…, z*_*n*_)^*⊤*^ is introduced, where *z*_*i*_ = *k* if the *i*-th sample point belongs to cluster *k* for *k* = 1, 2 *…, K*, where *K* is the number of clusters. *μ*_*kj*_ is the normalized expression level of gene *j* with *γ*_*j*_ = 1 observed in cluster *k*, while for a non-discriminating gene (i.e., *γ*_*j*_ = 0), we assume that gene *j* has the same normalized expression level *μ*_0*j*_ on all clusters. We assume a gamma prior for *μ*_*kj*_ and *μ*_0*j*_, that is, *μ*_*kj*_ ∼ Ga(*a*_*μ*_, *b*_*μ*_) and *μ*_0*j*_ ∼ Ga(*a*_0_, *b*_0_), and recommend small values such as *a*_*μ*_ = *b*_*μ*_ = *a*_0_ = *b*_0_ = 0.001 for a weakly informative setting.

### 2.3 Integrating the SRT Geospatial Profile via an MRF Prior Model

To efficiently incorporate available spatial information, we impose a Markov random field (MRF) prior on ***z*** to encourage neighboring samples to be clustered into the same group:

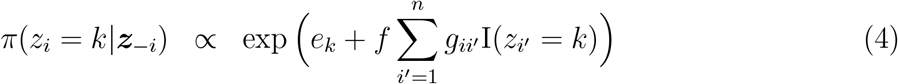

where *e*_*k*_ and *f* are hyperparameters to be chosen and ***z***_-*i*_ denotes the vector of ***z*** excluding the *i*-th element. Here *e*_*k*_ controls the abundance of each cluster, and *f* controls the strength of spatial dependency. As a result, neighboring samples are more likely to be assigned to the same cluster. The adjacency matrix ***G***, an *n × n* symmetric matrix, can be constructed based on the geospatial profile ***T*** to define the neighborhood structure, with *g*_*ii*_*′* = 1 if samples *I* and *i*^*′*^ are neighbors, and *g*_*ii*_*′* = 0 otherwise. For ST and 10x Visium platforms, ***G*** can be created based on their square and triangular lattices respectively. For other SRT technologies, we construct ***G*** using a Voronoi diagram (Boots et al., 2009) (see an example in Figure 4a). Note that if a sample point does not have any neighbors, its prior distribution reduces to a non-informative prior with parameter exp 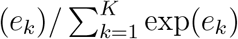 which is a multinomial logistic transformation of *e*_*k*_. Although the parameterization is somewhat arbitrary, careful consideration is required to determine the value of *f*. In particular, a large value of *f* may lead to a phase transition problem (i.e., all sample points are assigned into the same cluster). This problem arises because Equation (4) can only increase as a function of the number of I(*z*_*i*_ = *k*)’s equal to 1. In this paper, we regard *e* and *f* as fixed hyperparameters, following work by Li and Zhang (2010).

## 3. Model Fitting

### 3.1 MCMC Algorithm

We use a Markov chain Monte Carlo (MCMC) algorithm to update all parameters in our model. Our model allows simultaneously partitioning sample points into distinct clusters *via* ***z*** and identifying discriminating genes through ***γ***. We update the cluster allocation parameters *z*_*i*_’s and false zero indicators *η*_*ij*_’s using a Gibbs sampler. We jointly update the discriminating genes indicator *γ*_*j*_’s, normalized expression levels 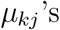 and 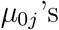 via an *add-delete* algorithm, and update the remaining parameters using the random walk Metropolis-Hastings (RWMH) algorithm. We note that this algorithm is sufficient to guarantee ergodicity for our model. For more details, please refer to the Supplementary Materials.

### 3.2 Posterior Inference

Our primary goal is to identify discriminating genes via the vector ***γ*** and cluster samples via the vector ***z***. We obtain posterior inference on these parameters by postprocessing the MCMC samples after burn-in. A common way to summarize the posterior distribution of ***z*** is by utilizing the *maximum a posteriori* (MAP) estimates, which can be calculated as follows:

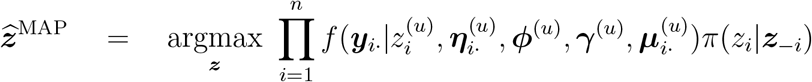

where *u* = 1,…, *U* indicates the MCMC iterations after burn-in. Apart from the MAP estimates, we may also obtain a summary of ***z*** based on the pairwise probability matrix (PPM). The PPM is an *n*-by-*n* symmetric matrix whose elements are the posterior pairwise probabilities of co-clustering, that is, the probability that sample *i* and sample *i′* are assigned to the same cluster: 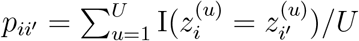 (Dahl, 2006). Then, the point estimate of the clustering, 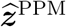, can be obtained by minimizing the sum of squared deviations of its association matrix from the PPM:

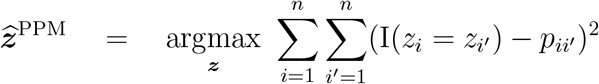

The PPM estimate has the advantage of utilizing information from all clusterings through the PPM. It is also intuitively appealing because it selects the “average” clustering rather than forming a clustering via an external, *ad hoc* clustering algorithm.

For feature selection, we can compute the MAP estimate of ***γ*** as follows:

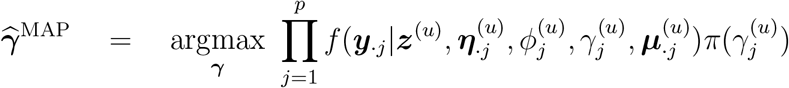

A more comprehensive way to summarize ***γ*** is based on their marginal posterior probabilities of inclusion (PPI), where 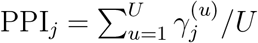 Then, the discriminating features are identified if their PPI values exceed a given threshold *c*:

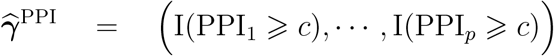

We can use the threshold *c* = 0.5, which is commonly referred to as the median model.

Alternatively, we can determine the threshold that controls for multiplicity (Newton et al., 2004), which ensures that the expected Bayesian false discovery rate (BFDR) is less than a specified value. The BFDR is computed as follows:

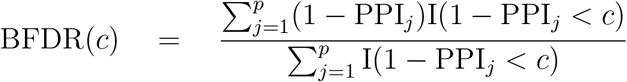

where BFDR(*c*) is the desired significance level.

## 4. Simulation Study

We used simulated data to evaluate the performance of our model and compare it with competing methods. We followed the same data generative schemes described in Li et al. (2021) and Jiang et al. (2022) based on two real spatial patterns respectively constructed from a mouse olfactory bulb (MOB) study and a human breast cancer (BC) study, which are depicted in Figure 2a. The MOB pattern and BC pattern respectively contain *n* = 260 and 250 spots. We simulated *p* = 100 genes, with *p*_*γ*_ = 15 discriminating genes.

**Figure 2:**
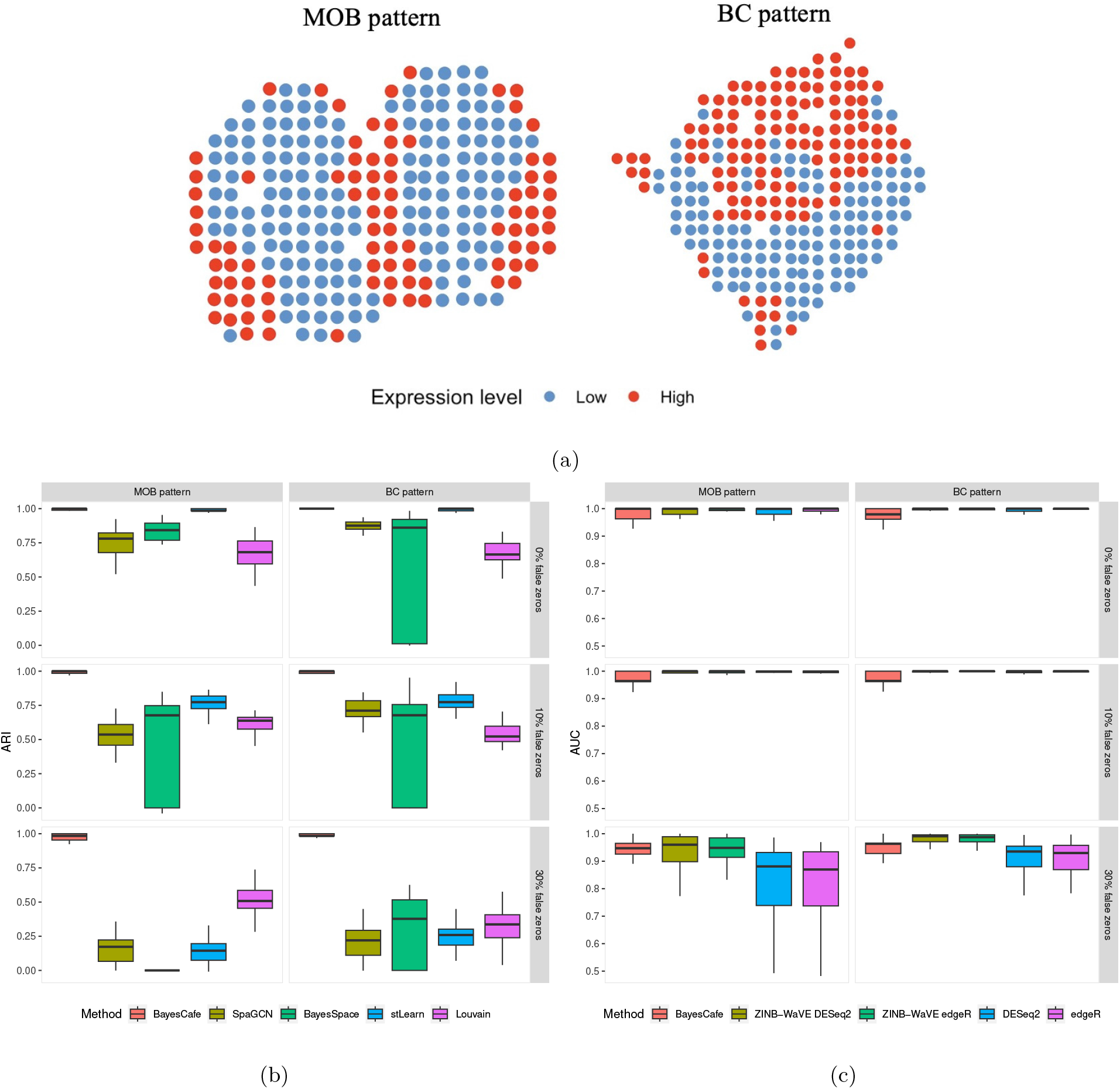
The simulation study. (a) The two spatial patterns used to generate the simulated data, which were constructed from the mouse olfactory bulb (MOB) and human breast cancer (BC) study, respectively. (b) The boxplots of adjusted Rand indices (ARIs) achieved by BayesCafe, SpaGCN, BayesSpace, stLearn, and Louvain under different scenarios in terms of spatial pattern and sparsity setting. (c) The boxplots of area under curves (AUCs) achieved by BayesCafe, ZINB-WaVE DESeq2, ZINB-WaVE edgeR, DESeq2, and edgeR under different scenarios in terms of spatial pattern and sparsity setting.

For each gene *j* at spot *i*, its latent normalized expression level was generated as:

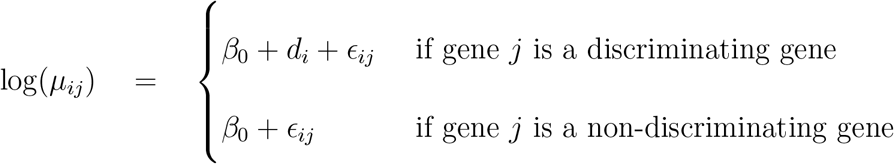

We set *β*_0_ = 2 as the baseline of log-normalized expression levels and ϵ_*ij*_ ∼ N(0, 0.3^2^) as the non-spatial random error. The spatial patterns were embedded by constructing *d*_*i*_’s. Therefore, for a non-discriminating gene, the normalized expression level log(*μ*_*ij*_) was i.i.d. from LN(2, 0.3^2^), resulting in no observed spatial correlation. For a discriminating gene with real patterns, each sample was dichotomized into two groups: high and low expression levels with *d*_*i*_ = log 3 and 0, respectively; therefore, the difference in *d*_*i*_’s between the two groups of spots introduced spatial differential expression patterns. To characterize the excess zeros and over-dispersion in the SRT data, we simulated the expression count *y*_*ij*_ from a ZINB distribution as follows:

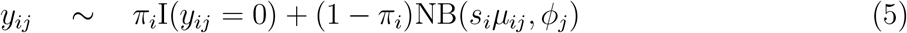

The size factor *s*_*i*_’s were i.i.d. from LN(0, 0.2^2^) and the gene-specific dispersion parameter *ϕ*_*j*_’s were i.i.d from Exp(0.1). For the choice of extra zero proportion *π*_*i*_, we randomly selected 0%, 10%, and 30% counts and manually set their values to zero. Combined with the two patterns and three zero-inflation settings, there were six different scenarios in total. For each scenario, we independently repeated the above steps to generate 30 replicates.

For the hyperparameter settings, we used the following defaults. We set the hyperparameters that control the proportion of extra zeros *a priori* to *π*_*i*_ ∼ Be(*a*_*π*_ = 1, *b*_*π*_ = 1). For the Beta prior on the feature selection parameter *ω*, we set *a*_*ω*_ = 0.1 and *b*_*ω*_ = 1.9, resulting in the proportion of features expected *a priori* to be *a*_*ω*_*/*(*a*_*ω*_ +*b*_*ω*_) = 5%. For the gamma priors on the NB dispersion parameter and normalized expression levels parameters, i.e. *ϕ*_*j*_ ∼ Ga(*a*_*ϕ*_, *b*_*ϕ*_), *μ*_0*j*_ ∼ Ga(*a*_0_, *b*_0_), and *μ*_*kj*_ ∼ Ga(*a*_*μ*_, *b*_*μ*_), we set *a*_*ϕ*_, *b*_*ϕ*_, *a*_0_, *b*_0_, *a*_*μ*_, and *b*_*μ*_ to 0.001, which led to a vague distribution with mean and variance respectively equal to 1 and 1000. For the MRF prior, we set *e*_1_ = *e*_2_ = 1 and *f* = 1. The results we report below were obtained by running one MCMC chain with 5000 iterations, discarding the first 50% of sweeps as burn-in, satisfying good MCMC convergence (Figure S2). We started the chain from a model with all *γ*_*j*_’s set to 0 and with each sample randomly assigned to a cluster with equal probability. All experiments were implemented in `R` with the `Rcpp` package to accelerate computations.

To compare clustering performance, we used the adjusted Rand index (ARI) (Hubert and Arabie, 1985), a corrected-for-chance version of the Rand index (Rand, 1971). The Rand index is used to measure the similarity between two different partitions and ranges between 0 and 1. Larger ARI values indicate better clustering results, and a value of 1 indicates a perfect match between two partitions.

To assess the performance of identifying the discriminating genes via the binary vector ***γ***, we utilized the area under the curve (AUC) of the receiver operating characteristic, a widely used metric in the evaluation of binary classifiers. AUC considers both the true positive rate (TPR) and false positive rate (FPR) at various threshold settings and ranges from 0 to 1; the higher the value, the more accurate the results.

Figure 2b displays the boxplots of ARIs achieved by different methods over 30 replicates under the six scenarios. As shown in the figure, all methods achieved good performance with no false zeros on the simulated data. BayesCafe and stLearn had similar performances and outperformed other methods under this setting. When the proportion of false zeros increased, BayesCafe clearly outperformed other methods and maintained nearly unaffected performance, implying that realistic modeling (i.e., accounting for zero-inflation) delivered an advantage over other methods. In contrast, all other methods suffered from decreased power, suggesting that the variance caused by an excess of zero counts was not properly addressed. To evaluate the performance of identifying discriminating genes, we compared BayesCafe with alternative methods, we used the results of stLearn clustering as input for clusters that produced better results than competing clustering methods. The competitor pool included DESeq2 (Love et al., 2014), edgeR (Robinson et al., 2010), ZINB-WaVE DESeq2 (Risso et al., 2018), and ZINB-WaVE edgeR (Risso et al., 2018), which are widely used to identify differentially-expressed (DE) genes in single-cell RNA-sequencing (scRNA-seq) data. The latter two methods are the modified versions of edgeR and DESeq2, using a ZINB-based Wanted Variation Extraction (ZINB-WaVE) strategy to downweight the inflated number of zeros in scRNA-seq data. edgeR uses an exact binomial test generalized for over-dispersed counts, while DESeq2 employs a Wald test by adopting a generalized linear model based on an NB kernel. Figure 2c displays the boxplots of AUCs achieved by different methods under different scenarios in terms of spatial pattern and sparsity settings. We observed that all methods achieved high performance with a low proportion of false zeros (i.e., 0% and 10% false zeros), with BayesCafe obtaining average AUCs over 0.96 under these two sparsity settings. Under the high sparsity setting (i.e., 30%), DESeq2 and edgeR experienced decreased power, indicating they cannot properly handle the zero-inflation problem. On the other hand, BayesCafe, ZINB-WaVE DESeq2, and ZINB-WaVE edgeR maintained high accuracy in detecting discriminating genes. BayesCafe achieved higher performance than all other methods for the BC pattern with an average AUC over 0.95, and achieved stable performance for the MOB pattern with an average AUC of 0.92. As shown in Figure S2, the discriminating genes identified by BayesCafe exhibit the same ability as truly discriminating genes from the PCA to explain the variance, indicating high identification accuracy.

In all, the simulation study clearly demonstrated that the joint modeling of spatial cluster structure and the associated discriminating genes via BayesCafe can boost the performance of both tasks.

## 5. Real Data Analysis

### 5.1 Application to the Mouse Olfactory Bulb ST Data

To further evaluate BayesCafe’s performance, we first examined a publicly available ST dataset from a mouse olfactory bulb (MOB) study (Ståhl et al., 2016). It is accessible on the website of the Spatial Research Lab at the KTH Royal Institute of Technology https://www.spatialresearch.org/. There are 12 replicates; we used replicate 12, which contains 16034 genes measured on 282 spots. The MOB data includes four main anatomic layers (i.e., clusters) organized in an inside–out fashion, annotated by CARD (Ma and Zhou, 2022) based on histology: the granule cell layer (GCL), the mitral cell layer (MCL), the glomerular layer (GL) and the nerve layer (ONL) (Figure 3a). We filtered out spots with fewer than 100 total counts across all genes and genes with more than 90% zero read counts on all spots and a maximum count of less than 10. This quality control procedure led to a final set of *n* = 278 spots and *p* = 1117 genes.

**Figure 3:**
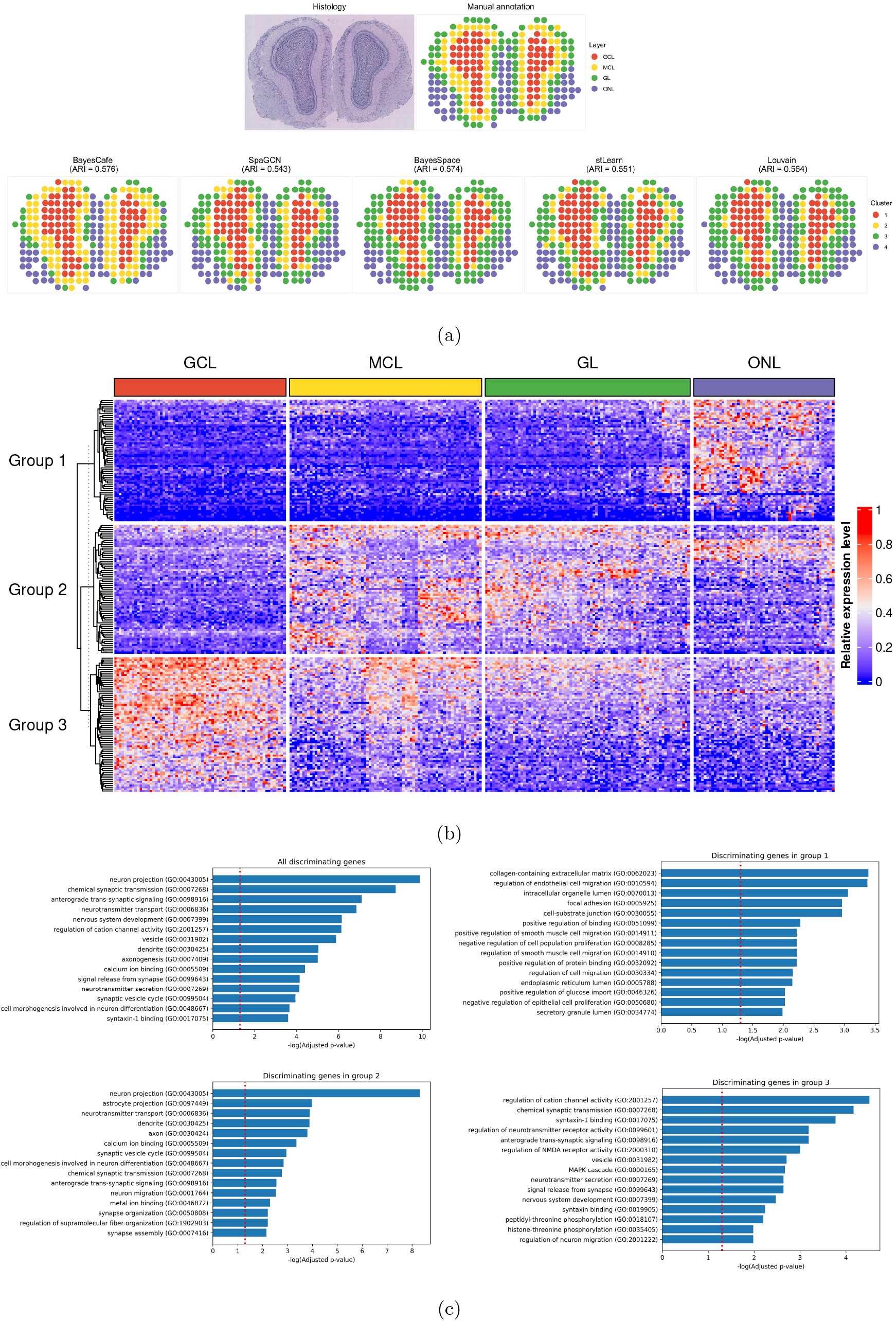
The mouse olfactory bulb (MOB) data analysis. (a) The hematoxylin and eosin (H&E)-stained tissue slide of the analyzed MOB ST data, with manual annotation serving as ground truth, and clusters detected by BayesCafe, SpaGCN, BayesSpace, stLearn, and Louvain. (b) The heatmap of discriminating genes across different layers detected by BayesCafe. (c) The Gene Ontology (GO) enrichment analysis of the discriminating genes identified by BayesCafe, with the red dashed line indicating a significance level of 0.05.

We evaluated the convergence of the MCMC algorithm based on the PPI vector of ***γ*** to calculate the PPIs for all four chains and found that their pairwise Pearson correlation coefficients ranged from 0.96 to 0.98, indicating good MCMC convergence. Figure S3 displays the trace plots of ARIs and number of discriminating genes across iterations in four chains; both of them indicate good MCMC convergence. We then aggregated the outputs of all four chains. We compared the clustering result of BayesCafe with SpaGCN, BayesSpace, stLearn, and Louvain, using default settings, and set the number of clusters as four for all methods. As Figure 3a shows that BayesCafe (ARI = 0.576) and BayesSpace (ARI = 0.574) achieved better performance while SpaGCN (ARI = 0.543), stLearn (ARI = 0.551), and Louvain (ARI = 0.564) generated inferior results. All methods were able to distinguish GCL and ONL layers well, but compared to BayesCafe, the other methods created a blurrier boundary between MCL and GL, resulting in inferior performance.

Next, we examined the discriminating genes detected by BayesCafe. Figure S3 shows the estimated marginal PPIs, Pr(*γ*_*j*_ = 1|*·*), of each single gene after burn-in. A threshold of 0.5 on the marginal probabilities results in a median model that includes 206 genes. To demonstrate the importance of these discriminating genes, we performed principal components analysis (PCA). We first normalized the counts in each sample (i.e., *y*_*ij*_*/s*_*i*_), and then rescaled each gene to ensure it had zero-mean and unit-variance. As shown in Figure S3, the discriminating genes could explain more variations for the first 50 PCs (around 25% of total discriminating genes). Specifically, the first 50 PCs of discriminating genes can explain about 78% of total variations of the data, whereas those of all 1117 genes after the quality control procedure can explain only around 46%. Moreover, the first 50 PCs of 100 sets, each containing 206 genes randomly chosen from all 1117 genes, obtained an average cumulative proportion of 64%. This comparison clearly shows that the 206 identified discriminating genes have a more significant impact on separating spots.

In addition, we examined the differential expression of discriminating genes among layers using the `Heatmap()` function from the `ComplexHeatmap R` package. The heatmap in Figure 3b shows three distinct gene groups that represent the major expression patterns observed in the data. Discriminating genes in group 1 showed enriched expression patterns in ONL, while those in group 2 expressed highly in layers other than GCL. Discriminating genes in the last group had a high expression level in GCL. The similar expression patterns between MCL and GL could explain why BayesCafe was unable to distinguish these two layers very well. This analysis validates the hypotheses that discriminating genes identified by BayesCafe express differentially among layers, and that BayesCafe could lead to biologically meaningful clusters. We can treat these identified discriminating genes as spatially variable (SV) genes as their expressions display spatially correlated patterns (Sun et al., 2020; Svensson et al., 2018).

To further demonstrate BayesCafe-defined discriminating genes align well with known biological knowledge, we compared SV genes detected by BayesCafe, SPARK (Sun et al., 2020), and SpatialDE (Svensson et al., 2018), with known olfactory bulb-specific gene set defined in Harmonizome database (PMID: 27374120). We found that the SV genes detected by BayesCafe showed higher overlap with known olfactory bulb-specific gene set than the ones defined by SPARK or SpatialDE (Figure S3), indicating that BayesCafe is able to identify biologically meaningful gene sets when analyzing spatial transcriptomics datasets.

Finally, we performed the gene ontology (GO) enrichment analysis using a Python wrapper GSEAPY (Kuleshov et al., 2016; Subramanian et al., 2007) on the selected discriminating genes to explore their relevant biological functions. Since genes in different groups show enriched expression patterns in different layers, we also performed the GO enrichment analysis of discriminating genes in three groups respectively. Table 1 presents a brief summary of the GO enrichment analysis results. GO enrichment analysis using all discriminating genes reported that a total of 2397 mouse GO terms in three components (biological processes, cellular components, and molecular functions) had at least one gene overlap with detected discriminating gene pools. Using an adjusted *p*-value smaller than 0.05 as the threshold, we found 193 enriched biological process GO terms. The top 15 GO terms with the smallest *p*-values are shown in Figure 3c. The figure indicates that many of the synaptic signaling and the nervous system-related GO terms were significantly enriched, both of which play important roles in synaptic organization and nerve development (Treloar et al., 2010). This indicates that the functions of those detected discriminating genes were associated with the underlying mechanism that drove the differentially expressed pattern. The individual GO enrichment analysis showed that discriminating genes in group 3 had more related enriched GO terms, and most of them were associated with biological processes, while discriminating genes in another two groups had a higher proportion of enriched GO terms related to cellular components (Figure 3c). This part strongly supports the hypothesis that identifying discriminating genes could uncover the underlying biological processes or functions.

**Table 1:**
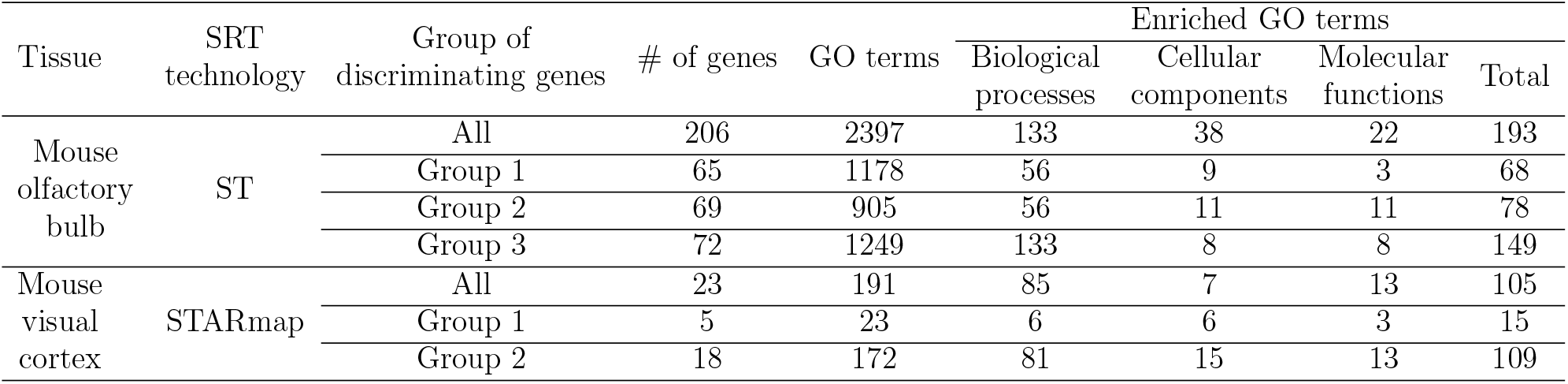
A summary of Gene Ontology (GO) enrichment analysis results of the mouse olfactory bulb (MOB) and mouse visual cortex data.

| Tissue | SRT technology | Group of discriminating genes | # of genes | GO terms | Enriched GO terms |  |  | Total |
| --- | --- | --- | --- | --- | --- | --- | --- | --- |
|  |  |  |  |  | Biological processes | Cellular components | Molecular functions |  |
| Mouse olfactory bulb | ST | All | 206 | 2397 | 133 | 38 | 22 | 193 |
|  |  | Group 1 | 65 | 1178 | 56 | 9 | 3 | 68 |
|  |  | Group 2 | 69 | 905 | 56 | 11 | 11 | 78 |
|  |  | Group 3 | 72 | 1249 | 133 | 8 | 8 | 149 |
| Mouse visual cortex | STARmap | All | 23 | 191 | 85 | 7 | 13 | 105 |
|  |  | Group 1 | 5 | 23 | 6 | 6 | 3 | 15 |
|  |  | Group 2 | 18 | 172 | 81 | 15 | 13 | 109 |

### 5.2 Application to the Mouse Visual Cortex STARmap Data

We also applied BayesCafe to the mouse visual cortex STARmap dataset fromhttps://www.starmapresources.com/data at single-cell resolution (Wang et al., 2018). The STARmap dataset measured 1020 genes in 1207 cells, corresponding to 7 layers (Figure 4a). We used the same analysis procedures as described in Section 5.1, with *n* = 1207 cells and *p* = 105 genes remaining after our quality control procedure. The pairwise Pearson correlation coefficients of PPIs range from 0.7 to 0.77 for four MCMC chains, along with trace plots of ARIs and the number of discriminating genes in Figure S4, suggesting reasonable convergence.

**Figure 4:**
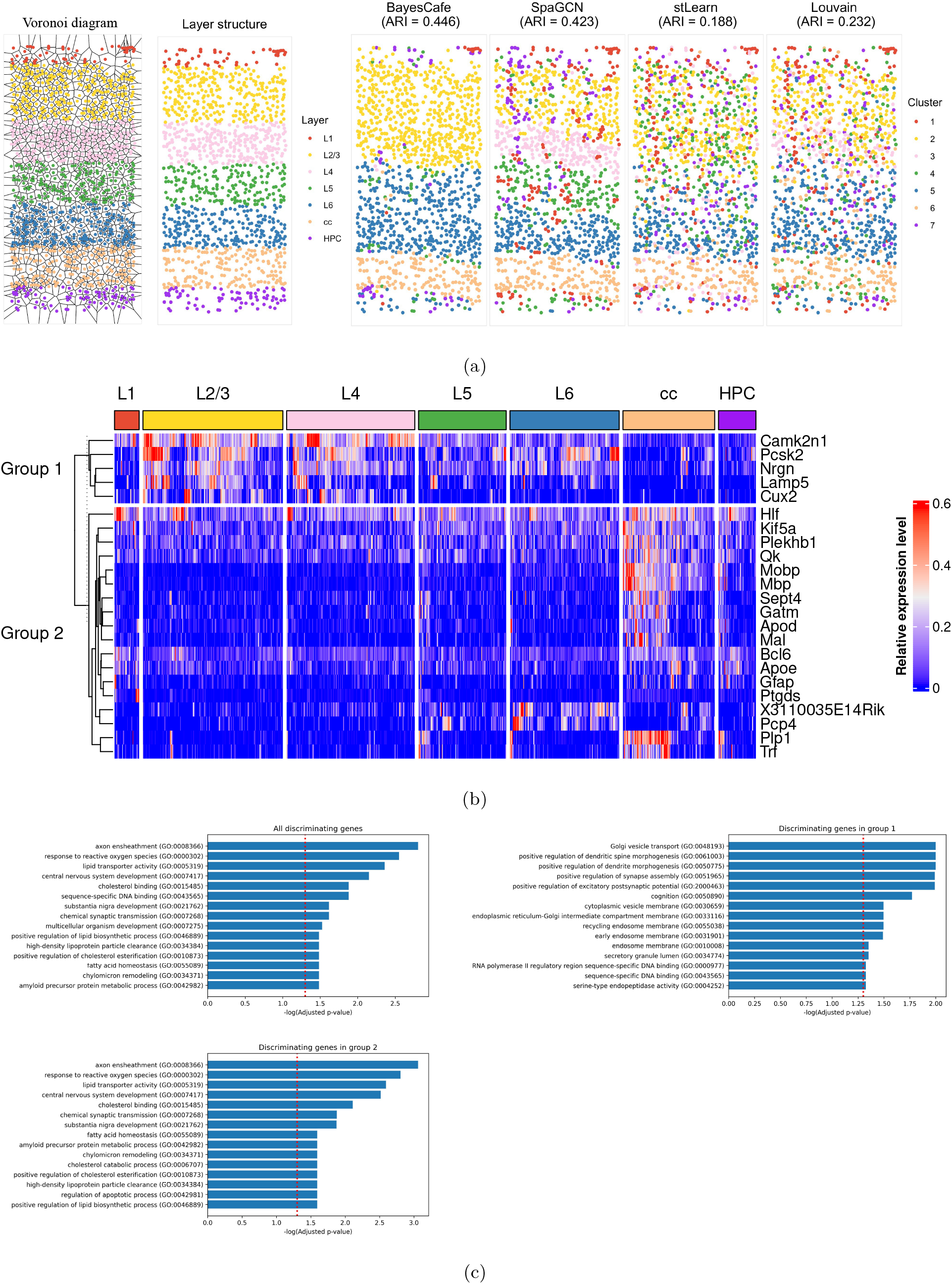
The mouse visual cortex data analysis. (a) The Voronoi diagram constructed for defining the neighboring structure, manual annotation served as ground truth, and clusters detected by BayesCafe, SpaGCN, stLearn, and Louvain. (b) The heatmap of discriminating genes across different layers detected by BayesCafe. (c) The Gene Ontology (GO) enrichment analysis of the discriminating genes identified by BayesCafe, with the red dashed line indicating a significance level of 0.05.

The neighboring structure was defined with a Voronoi diagram (Figure 4a), where two samples sharing the same edge were defined as neighbors. Figure 4a shows that BayesCafe outperformed all other methods by achieving the highest ARI of 0.446, but was unable to distinguish the L2/3 from L4 and L5 from L6, respectively. By contrast, the ARIs of other methods were much lower (0.423 for SpaGCN, 0.188 for stLearn, and 0.232 for Louvain). This dataset demonstrates that BayesCafe is able to incorporate spatial information more efficiently than SpaGCN and stLearn, and that the usage of feature selection could improve clustering accuracy.

BayesCafe detected 23 discriminating genes out of 105 genes with a threshold of 0.5 (Figure S4). Principal components analysis validated the hypothesis that identifying discriminating genes could greatly improve the separation of samples. As shown in Figure S4, the first five PCs of the discriminating genes can explain about 52% of the total variations, while those of all 105 genes after the quality control procedure only can explain around 23% of the total variations. In contrast, the first 5 PCs of 100 sets, each containing 23 genes randomly selected from all 105 genes, obtained an average cumulative proportion of 37%. As Figure 4b shows that discriminating genes were divided into two groups with the number of genes of 5 and 18, respectively. Discriminating genes in group 1 showed high expression levels in layers L2/3 and L4, while those in group 2 expressed highly in layers cc and HPC. We also point out that gene expression patterns are similar between L2/3 and L4, and between L5 and L6. This could be the reason that cells in layers L2/3 and L4, and L5 and L6, were clustered together in BayesCafe.

To further validate biological significance of clusters defined by BayesCafe, we extracted known visual cortex gene sets from the Allen Mouse Brain Atlas (PMID: 23193282) and compared with SV genes defined by BayesCafe, SPARK, and SpatialDE in the visual cortex samples based on STARMap technology. We found that BayesCafe outperformed SPARK or SpatialDE with higher enrichment of known visual cortex genes (Figure S4), confirming that BayesCafe-identified discriminating genes are more consistent with known biological knowledge than SPARK and SpatialDE.

Finally, we performed GO enrichment analysis using all discriminating genes as well as discriminating genes in group 1 and 2. As shown in Table 1, GO enrichment analysis using all discriminating genes reported a total of 191 GO terms with at least one overlapping gene. We found 105 enriched GO terms, with 85, 7, and 13 enriched GO terms related to biological processes, cellular components, and molecular functions, respectively. The individual GO enrichment analysis showed that discriminating genes in group 2 had more related enriched GO terms, and most of them were associated with biological processes, while 5 discriminating genes in group 1 still detected 15 enriched GO terms. Overall, these discoveries highlight the meaningful biological interpretations that can be inferred from the differentially-expressed discriminating genes.

## 6. Discussion

In this paper, we developed BayesCafe, a novel Bayesian ZINB mixture model that can account for the spatial correlation of SRT data by employing a MRF prior for clustering analysis and uses a feature selection approach to detect discriminating genes. Compared to existing methods, BayesCafe offers several advantages. Firstly, it directly models count data with an NB distribution, which can better account for over-dispersion compared to a Poisson distribution, providing a more accurate representation of the data. Secondly, it properly addresses the issue of excess zero counts commonly observed in SRT data by incorporating a ZINB model, resulting in more robust performance. Thirdly, it utilizes a feature selection mechanism that not only generates low-dimensional summaries of the SRT data, it also identifies the most discriminating genes, thus improving model performance and interpretability. Fourthly, it efficiently incorporates spatial information and obtains a more robust and accurate clustering result via the MRF prior model. Finally, it improves parameter estimations and uncertainties quantification using a Bayesian approach, leading to more reliable and quantifiable inference. In our simulation study, BayesCafe outperformed all other clustering methods, especially when a high number of zeros were present in the data. Furthermore, BayesCafe was able to detect the discriminating genes with higher and more stable accuracy. In our real data analysis, BayesCafe demonstrated higher accuracy in clustering analysis by incorporating spatial information and employing the feature selection procedure. In addition, discriminating genes identified by BayesCafe were differentially-expressed and associated with biological functions.

Several extensions of our model are worth investigating. Firstly, as it follows a Bayesian framework, BayesCafe can be extended to estimate the number of clusters *K* via the Dirichlet process mixture (Müller et al., 2015; Li et al., 2017). Secondly, BayesCafe can enable joint estimation of discriminating genes by incorporating pathway information as prior knowledge to consider regulatory relationships among genes (Li et al., 2019).

## Supporting information

Supplementary Materials for BayesCafe

## Acknowledgements

The authors would like to thank the chief editor, guest editor, and reviewer for their suggestions and feedback, which improved the paper significantly, and Kevin W. Jin for helping us proofread the manuscript.

## Supplementary Materials

Web Appendices, Table, and Figures referenced in Sections 2, 3, 4, and 5 are available with this paper at the Biometrics website on Wiley Online Library. All simulated and real data used for analysis, and the related source code in `R/C++` are available athttps://github.com/huimin230/BayesCafe.

## References

Blondel, V. D., Guillaume, J.-L., Lambiotte, R., and Lefebvre, E. (2008). Fast unfolding of communities in large networks. Journal of statistical mechanics: theory and experiment 2008, P10008.

Boots, B., Sugihara, K., Chiu, S. N., and Okabe, A. (2009). Spatial tessellations: concepts and applications of Voronoi diagrams.

Dahl, D. B. (2006). Model-based clustering for expression data via a Dirichlet process mixture model. Bayesian inference for gene expression and proteomics 4, 201–218.

De Bruin, E. C., McGranahan, N., Mitter, R., Salm, M., Wedge, D. C., Yates, L., Jamal-Hanjani, M., Shafi, S., Murugaesu, N., Rowan, A. J., et al. (2014). Spatial and temporal diversity in genomic instability processes defines lung cancer evolution. Science 346, 251–256.

Eng, C.-H. L., Lawson, M., Zhu, Q., Dries, R., Koulena, N., Takei, Y., Yun, J., Cronin, C., Karp, C., Yuan, G.-C., et al. (2019). Transcriptome-scale super-resolved imaging in tissues by RNA seqFISH+. Nature 568, 235–239.

Finak, G., McDavid, A., Yajima, M., Deng, J., Gersuk, V., Shalek, A. K., Slichter, C. K., Miller, H. W., McElrath, M. J., Prlic, M., et al. (2015). MAST: a flexible statistical framework for assessing transcriptional changes and characterizing heterogeneity in single-cell RNA sequencing data. Genome biology 16, 1–13.

Hu, J., Li, X., Coleman, K., Schroeder, A., Ma, N., Irwin, D. J., Lee, E. B., Shinohara, R. T., and Li, M. (2021). SpaGCN: Integrating gene expression, spatial location and histology to identify spatial domains and spatially variable genes by graph convolutional network. Nature methods 18, 1342–1351.

Hubert, L. and Arabie, P. (1985). Comparing partitions. Journal of classification 2, 193–218.

Jiang, X., Xiao, G., and Li, Q. (2022). A Bayesian modified Ising model for identifying spatially variable genes from spatial transcriptomics data. Statistics in medicine 41, 4647–4665.

Kuleshov, M. V., Jones, M. R., Rouillard, A. D., Fernandez, N. F., Duan, Q., Wang, Z., Koplev, S., Jenkins, S. L., Jagodnik, K. M., Lachmann, A., et al. (2016). Enrichr: a comprehensive gene set enrichment analysis web server 2016 update. Nucleic acids research 44, W90–W97.

Li, F. and Zhang, N. R. (2010). Bayesian variable selection in structured high-dimensional covariate spaces with applications in genomics. Journal of the American statistical association 105, 1202–1214.

Li, Q., Cassese, A., Guindani, M., and Vannucci, M. (2019). Bayesian negative binomial mixture regression models for the analysis of sequence count and methylation data. Biometrics 75, 183–192.

Li, Q., Guindani, M., Reich, B. J., Bondell, H. D., and Vannucci, M. (2017). A Bayesian mixture model for clustering and selection of feature occurrence rates under mean constraints. Statistical analysis and data mining: The ASA data science journal 10, 393–409.

Li, Q., Zhang, M., Xie, Y., and Xiao, G. (2021). Bayesian modeling of spatial molecular profiling data via Gaussian process. Bioinformatics 37, 4129–4136.

Liu, W., Liao, X., Yang, Y., Lin, H., Yeong, J., Zhou, X., Shi, X., and Liu, J. (2022). Joint dimension reduction and clustering analysis of single-cell RNA-seq and spatial transcriptomics data. Nucleic acids research 50, e72–e72.

Love, M. I., Huber, W., and Anders, S. (2014). Moderated estimation of fold change and dispersion for RNA-seq data with DESeq2. Genome biology 15, 1–21.

Lubeck, E., Coskun, A. F., Zhiyentayev, T., Ahmad, M., and Cai, L. (2014). Single-cell insitu RNA profiling by sequential hybridization. Nature methods 11, 360–361.

Lun, A. T., Bach, K., and Marioni, J. C. (2016). Pooling across cells to normalize single-cell RNA sequencing data with many zero counts. Genome biology 17, 1–14.

Ma, Y. and Zhou, X. (2022). Spatially informed cell-type deconvolution for spatial transcriptomics. Nature biotechnology pages 1–11.

Marx, V. (2021). Method of the year: spatially resolved transcriptomics. Nature methods 18, 9–14.

McInnes, L., Healy, J., and Melville, J. (2018). Umap: Uniform manifold approximation and projection for dimension reduction. arXiv preprint 1802.03426.

Müller, P., Quintana, F. A., Jara, A., and Hanson, T. (2015). Bayesian nonparametric data analysis, volume 1. Springer.

Newton, M. A., Noueiry, A., Sarkar, D., and Ahlquist, P. (2004). Detecting differential gene expression with a semiparametric hierarchical mixture method. Biostatistics 5, 155–176.

Pham, D., Tan, X., Xu, J., Grice, L. F., Lam, P. Y., Raghubar, A., Vukovic, J., Ruitenberg, M. J., and Nguyen, Q. (2020). stLearn: integrating spatial location, tissue morphology and gene expression to find cell types, cell-cell interactions and spatial trajectories within undissociated tissues. BioRxiv pages 2020–05.

Rand, W. M. (1971). Objective criteria for the evaluation of clustering methods. Journal of the American statistical association 66, 846–850.

Risso, D., Perraudeau, F., Gribkova, S., Dudoit, S., and Vert, J.-P. (2018). A general and flexible method for signal extraction from single-cell RNA-seq data. Nature communications 9, 284.

Robinson, M. D., McCarthy, D. J., and Smyth, G. K. (2010). edgeR: a Bioconductor package for differential expression analysis of digital gene expression data. Bioinformatics 26, 139–140.

Rodriques, S. G., Stickels, R. R., Goeva, A., Martin, C. A., Murray, E., Vanderburg, C. R., Welch, J., Chen, L. M., Chen, F., and Macosko, E. Z. (2019). Slide-seq: A scalable technology for measuring genome-wide expression at high spatial resolution. Science 363, 1463–1467.

Satija, R., Farrell, J. A., Gennert, D., Schier, A. F., and Regev, A. (2015). Spatialreconstruction of single-cell gene expression data. Nature biotechnology 33, 495–502.

Ståhl, P. L., Salmén, F., Vickovic, S., Lundmark, A., Navarro, J. F., Magnusson, J., Giacomello, S., Asp, M., Westholm, J. O., Huss, M., et al. (2016). Visualization and analysis of gene expression in tissue sections by spatial transcriptomics. Science 353, 78–82.

Subramanian, A., Kuehn, H., Gould, J., Tamayo, P., and Mesirov, J. P. (2007). GSEA-P: a desktop application for Gene Set Enrichment Analysis. Bioinformatics 23, 3251–3253.

Sun, S., Zhu, J., and Zhou, X. (2020). Statistical analysis of spatial expression patterns for spatially resolved transcriptomic studies. Nature methods 17, 193–200.

Svensson, V., Teichmann, S. A., and Stegle, O. (2018). SpatialDE: identification of spatially variable genes. Nature methods 15, 343–346.

Tadesse, M. G., Sha, N., and Vannucci, M. (2005). Bayesian variable selection in clustering high-dimensional data. Journal of the American statistical association 100, 602–617.

Treloar, H. B., Miller, A. M., Ray, A., and Greer, C. A. (2010). Development of the olfactory system. The neurobiology of olfaction 20092457, 131–155.

Van der Maaten, L. and Hinton, G. (2008). Visualizing data using t-SNE. Journal of machine learning research 9,.

Wang, X., Allen, W. E., Wright, M. A., Sylwestrak, E. L., Samusik, N., Vesuna, S., Evans, K., Liu, C., Ramakrishnan, C., Liu, J., et al. (2018). Three-dimensional intact-tissue sequencing of single-cell transcriptional states. Science 361, eaat5691.

Yang, Y., Shi, X., Liu, W., Zhou, Q., Chan Lau, M., Chun Tatt Lim, J., Sun, L., Ng, C. C. Y., Yeong, J., and Liu, J. (2022). SC-MEB: spatial clustering with hidden markov random field using empirical Bayes. Briefings in bioinformatics 23, bbab466.

Zhao, E., Stone, M. R., Ren, X., Guenthoer, J., Smythe, K. S., Pulliam, T., Williams, S. R., Uytingco, C. R., Taylor, S. E., Nghiem, P., et al. (2021). Spatial transcriptomics at subspot resolution with BayesSpace. Nature biotechnology 39, 1375–1384.

