## Supplementary Materials for BayesCafe for "An Interpretable Bayesian Clustering Approach with Feature Selection for Analyzing Spatially Resolved Transcriptomics Data"

### 1. Full Details of the MCMC Algorithm

The model parameter space consists of $(\boldsymbol{z}, \boldsymbol{H}, \boldsymbol{\phi}, \boldsymbol{\gamma}, \boldsymbol{M})$, where $\boldsymbol{z} = \{z_i, i = 1, \cdots, n\}$ is the cluster allocation vector, $\boldsymbol{H} = \{\eta_{ij}, i = 1, \cdots, n, j = 1, \cdots, p\}$ is the extra zero indicator matrix, $\boldsymbol{\phi} = \{\phi_j, j = 1, \cdots, p\}$ is the collection of dispersion parameters, $\boldsymbol{\gamma} = \{\gamma_j, j = 1, \cdots, p\}$ is the discriminating gene indicator vector, and $\boldsymbol{M} = \{u_{kj}, k = 1, \cdots, K, j = 1, \cdots, p\}$ is the collection of normalized expression levels, which can be divided into 1) $\boldsymbol{\mu}_0 = \{\mu_{0j}, j : \gamma_j = 0\}$ and $\boldsymbol{M}^{(\gamma)} = \{u_{kj}, k = 1, \cdots, K, j : \gamma_j = 1\}$. We start by writing the full posterior:

$$\pi(\boldsymbol{z}, \boldsymbol{H}, \boldsymbol{\phi}, \boldsymbol{\gamma}, \boldsymbol{M}|\boldsymbol{Y}) \quad \propto \quad f(\boldsymbol{Y}|\boldsymbol{z}, \boldsymbol{H}, \boldsymbol{\phi}, \boldsymbol{\gamma}, \boldsymbol{M})\pi(\boldsymbol{z})\pi(\boldsymbol{H})\pi(\boldsymbol{\phi})\pi(\boldsymbol{\gamma})\pi(\boldsymbol{M})$$

According to Section 2.1, we can compute the data likelihood and priors as

$$\begin{aligned}
f(\boldsymbol{Y}|\boldsymbol{z}, \boldsymbol{H}, \boldsymbol{\phi}, \boldsymbol{\gamma}, \boldsymbol{M}) &= \prod_{i=1}^{n} \left( \prod_{\{j:\eta_{ij}=0,\gamma_j=1\}} \mathrm{NB}(y_{ij}; s_i\mu_{kj}, \phi_j) \prod_{\{j:\eta_{ij}=0,\gamma_j=0\}} \mathrm{NB}(y_{ij}; s_i\mu_{0j}, \phi_j) \right) \\
\pi(\boldsymbol{H}) &= \prod_{i=1}^{n}\prod_{j=1}^{p} \text{Be-Bern}(\eta_{ij}; a_\pi, b_\pi) \\
\pi(\boldsymbol{\phi}) &= \prod_{j=1}^{p} \mathrm{Ga}(\phi_j; a_\phi, b_\phi)
\end{aligned}$$

and according to Sections 2.2 and 2.3, we can calculate the rest of priors as

$$\begin{aligned}
\pi(\boldsymbol{\gamma}) &= \prod_{j=1}^{p} \text{Be-Bern}(\gamma_j; a_\omega, b_\omega) \\
\pi(\boldsymbol{z}) &\propto \prod_{i=1}^{n} \exp\left( e_k + f\sum_{i'=1}^{n} g_{ii'}\mathrm{I}(z_{i'} = k) \right) \\
\pi(\boldsymbol{M}) &= \prod_{k=1}^{K} \left( \prod_{\{j:\gamma_j=1\}} \mathrm{Ga}(\mu_{kj}; a_\mu, b_\mu) \prod_{\{j:\gamma_j=0\}} \mathrm{Ga}(\mu_{0j}; a_\mu, b_\mu) \right)
\end{aligned}$$

The pmf's or pdf's of the involved common distributions are given below:

$$\text{If} \quad x \sim \mathrm{NB}(\mu, \phi), \quad \text{then} \quad p(x) = \frac{\Gamma(x+\phi)}{x!\Gamma(\phi)}\left(\frac{\phi}{\mu+\phi}\right)^{\phi}\left(\frac{\mu}{\mu+\phi}\right)^{x}$$

$$\text{If} \quad x \sim \text{Be-Bern}(a, b), \quad \text{then} \quad p(x) = \frac{1}{a+b}\frac{\Gamma(a+x)\Gamma(b+1-x)}{\Gamma(a)\Gamma(b)}$$

$$\text{If} \quad x \sim \mathrm{Ga}(a, b), \quad \text{then} \quad p(x) = \frac{b^a}{\Gamma(a)}x^{a-1}\exp(-bx)$$

We detail the MCMC algorithm below. At each MCMC iteration, We perform the following steps:

**Update of cluster allocation $\boldsymbol{z}$:** We update each $z_i, i = 1, \cdots, n$ separately using a Gibbs sampler. At each iteration, we draw a new value $z_i = k$ with the probability $\pi(z_i = k|\cdot)/\sum_{k'=1}^{K} \pi(z = k'|\cdot)$, where

$$\pi(z_i = k|\cdot) \quad \propto \quad \prod_{\{j:\eta_{ij}=0,\gamma_j=1\}} \mathrm{NB}(y_{ij}; s_i\mu_{kj}, \phi_j) \exp\left(e_k + f\sum_{i'=1}^{n} g_{ii'}\mathrm{I}(z_{i'} = k)\right)$$

**Update of zero-inflation indicator $\boldsymbol{H}$:** We update each extra zero indicator $\eta_{ij}, i = 1, \cdots, n, j = 1, \cdots, p$ separately that corresponds to $y_{ij} = 0$ using the Gibbs sampler:

$$\begin{aligned}
\pi(\eta_{ij}|\cdot) \quad &\propto \quad (\mathrm{NB}(y_{ij}; s_i\mu_{kj}, \phi_j))^{\gamma_j}(\mathrm{NB}(y_{ij}; s_i\mu_{0j}, \phi_j))^{1-\gamma_j}\text{Be-Bern}(\eta_{ij}; a_\pi, b_\pi) \\
\eta_{ij}|\cdot \quad &\sim \quad \mathrm{Bern}\left(\frac{\pi(\eta_{ij=1}|\cdot)}{\pi(\eta_{ij=1}|\cdot) + \pi(\eta_{ij=0}|\cdot)}\right)
\end{aligned}$$

**Joint update of informative gene indicator $\boldsymbol{\gamma}$ and normalized gene expression levels $\boldsymbol{M}$:** We perform a between-model step to update these parameters jointly since $\boldsymbol{M}$ depends on $\boldsymbol{\gamma}$. This is done *via* an *add-delete* algorithm. In this approach, a new candidate vector, say $\boldsymbol{\gamma}^*$, is generated by randomly choosing an entry of $\boldsymbol{\gamma}$, say $j$, and changing its value to $\gamma_j^* = 1 - \gamma_j$. Then, this proposed move is accepted with probability $\min(1, m_{\mathrm{MH}})$, where the Hastings ratio is

$$m_{\mathrm{MH}} \quad = \quad \frac{f(\boldsymbol{y}_{\cdot j}|\boldsymbol{z}, \boldsymbol{\eta}_{\cdot j}, \phi_j, \gamma_j, \mu_{0j}^*, \boldsymbol{\mu}_{\cdot j}^*)}{f(\boldsymbol{y}_{\cdot j}|\boldsymbol{z}, \boldsymbol{\eta}_{\cdot j}, \phi_j, \gamma_j, \mu_{0j}, \boldsymbol{\mu}_{\cdot j})} \frac{\pi(\mu_{0j}^*, \boldsymbol{\mu}_{\cdot j}^*|\gamma_j^*)}{\pi(\mu_{0j}, \boldsymbol{\mu}_{\cdot j}|\gamma_j)} \frac{\pi(\gamma_j^*)}{\pi(\gamma_j)} \frac{J(\mu_{0j}, \boldsymbol{\mu}_{\cdot j} \leftarrow \mu_{0j}^*, \boldsymbol{\mu}_{\cdot j}^*|\gamma_j \leftarrow \gamma_j^*)}{J(\mu_{0j}^*, \boldsymbol{\mu}_{\cdot j}^* \leftarrow \mu_{0j}, \boldsymbol{\mu}_{\cdot j}|\gamma_j^* \leftarrow \gamma_j)} \frac{J(\gamma_j \leftarrow \gamma_j^*)}{J(\gamma_j^* \leftarrow \gamma_j)}$$

while the last proposal density ratio $J(\gamma_j \leftarrow \gamma_j^*)/J(\gamma_j^* \leftarrow \gamma_j)$ equals one. We use $J(\cdot \leftarrow \cdot)$ to denote the proposal probability function for selected move. For the *add* step, we propose a new $\log\mu_{kj}^*$ from $\mathrm{N}(\log\theta_0, (10\tau_\mu)^2)$. For the *delete* step, we propose a new $\log\mu_{0j}^*$ from

$\mathrm{N}(\log\theta_0, \tau_\mu^2)$, where $\theta_0$ is either $\exp(2)$ or $\sum_{i=1}^n \tilde{\mu}_{ij}/n$ and $\tilde{\mu}_{ij} = y_{ij}/s_i$. Then,

$$\frac{\pi(\mu_{0j}^*, \boldsymbol{\mu}_{\cdot j}^*|\gamma_j^*)}{\pi(\mu_{0j}, \boldsymbol{\mu}_{\cdot j}|\gamma_j)} = \begin{cases} \prod_{k=1}^K \pi(\mu_{kj}^*)/\pi(\mu_{0j}) & \text{for } \textit{add step} \\ \pi(\mu_{0j}^*)/\prod_{k=1}^K \pi(\mu_{kj}) & \text{for } \textit{delete step} \end{cases}$$

$$\frac{J(\mu_{0j}, \boldsymbol{\mu}_{\cdot j} \leftarrow \mu_{0j}^*, \boldsymbol{\mu}_{\cdot j}^*|\gamma_j \leftarrow \gamma_j^*)}{J(\mu_{0j}^*, \boldsymbol{\mu}_{\cdot j}^* \leftarrow \mu_{0j}, \boldsymbol{\mu}_{\cdot j}|\gamma_j^* \leftarrow \gamma_j)} = \begin{cases} \frac{\mathrm{N}(\log\mu_{0j};\log\theta_0,\tau_\mu^2)}{\prod_{k=1}^K \mathrm{N}(\log\mu_{kj}^*;\log\theta_0,(10\tau_\mu)^2)} & \text{for } \textit{add step} \\ \frac{\prod_{k=1}^K \mathrm{N}(\log\mu_{kj};\log\theta_0,(10\tau_\mu)^2)}{\mathrm{N}(\log\mu_{0j}^*;\theta_0,\tau_\mu^2)} & \text{for } \textit{delete step} \end{cases}$$

**Update of normalized expression levels $\boldsymbol{M}^{(\gamma)}$:** We perform a within-model step to update each $\mu_{kj}, k = 1, \cdots, K$ of $\gamma_j = 1$ separately *via* the random walk Metropolis-Hastings (RWMH) algorithm. We first propose a new $\log\mu_{kj}^*$ from $\mathrm{N}(\log\mu_{kj}, \tau_\mu^2)$ and then accept the proposed value $\mu_{kj}^*$ with probability $\min(1,\, m_{\mathrm{MH}})$, where the Hastings ratio is:

$$m_{\mathrm{MH}} = \frac{\prod_{\{i:\eta_{ij}=0,z_i=k\}} \mathrm{NB}(y_{ij}; s_i\mu_{kj}^*, \phi_j)}{\prod_{\{i:\eta_{ij}=0,z_i=k\}} \mathrm{NB}(y_{ij}; s_i\mu_{kj}, \phi_j)} \frac{\mathrm{Ga}(\mu_{kj}^*; a_\mu, b_\mu)}{\mathrm{Ga}(\mu_{kj}; a_\mu, b_\mu)} \frac{J(\mu_{kj} \leftarrow \mu_{kj}^*)}{J(\mu_{kj}^* \leftarrow \mu_{kj})}$$

Note that the proposal density ratio equals one for this RWMH update.

**Update of normalized gene expression levels $\boldsymbol{\mu}_0$:** We perform a within-model step to update each $\mu_{0j}$ of $\gamma_j = 0$ separately *via* the RWMH algorithm. We first propose a new $\log\mu_{0j}^*$ from $\mathrm{N}(\log\mu_{0j}, \tau_\mu^2)$ and then accept the proposed value $\mu_{0j}^*$ with probability $\min(1,\, m_{\mathrm{MH}})$, where the Hastings ratio is

$$m_{\mathrm{MH}} = \frac{\prod_{\{i:\eta_{ij}=0\}} \mathrm{NB}(y_{ij}; s_i\mu_{0j}^*, \phi_j)}{\prod_{\{i:\eta_{ij}=0\}} \mathrm{NB}(y_{ij}; s_i\mu_{0j}, \phi_j)} \frac{\mathrm{Ga}(\mu_{0j}^*; a_0, b_0)}{\mathrm{Ga}(\mu_{0j}; a_0, b_0)} \frac{J(\mu_{0j} \leftarrow \mu_{0j}^*)}{J(\mu_{0j}^* \leftarrow \mu_{0j})}$$

Note that the proposal density ratio equals one for this RWMH update.

**Update of dispersion parameter $\boldsymbol{\phi}$:** We update each $\phi_j, j = 1, \cdots, p$ separately by using the RWMH algorithm. We first propose a new $\log\phi_j^*$ from $\mathrm{N}(\log\phi_j, \tau_\phi^2)$ and then accept the proposed value $\phi_j^*$ with probability $\min(1,\, m_{\mathrm{MH}})$, where the Hastings ratio is

$$m_{\mathrm{MH}} = \frac{\left(\prod_{k=1}^K \prod_{\{i:\eta_{ij}=0,z_i=k\}} \mathrm{NB}(y_{ij}; s_i\mu_{kj}, \phi_j^*)\right)^{\gamma_j} \left(\prod_{\{i:\eta_{ij}=0\}} \mathrm{NB}(y_{ij}; s_i\mu_{0j}, \phi_j^*)\right)^{1-\gamma_j}}{\left(\prod_{k=1}^K \prod_{\{i:\eta_{ij}=0,z_i=k\}} \mathrm{NB}(y_{ij}; s_i\mu_{kj}, \phi_j)\right)^{\gamma_j} \left(\prod_{\{i:\eta_{ij}=0\}} \mathrm{NB}(y_{ij}; s_i\mu_{0j}, \phi_j)\right)^{1-\gamma_j}} \frac{\mathrm{Ga}(\phi_j^*; a_\phi, b_\phi)}{\mathrm{Ga}(\phi_j; a_\phi, b_\phi)} \frac{J(\phi_j \leftarrow \phi_j^*)}{J(\phi_j^* \leftarrow \phi_j)}$$

Note that the proposal density ratio equals one for this RWMH update.

[Figure 1 about here.]

[Figure 2 about here.]

[Figure 3 about here.]

[Figure 4 about here.]

[Table 1 about here.]

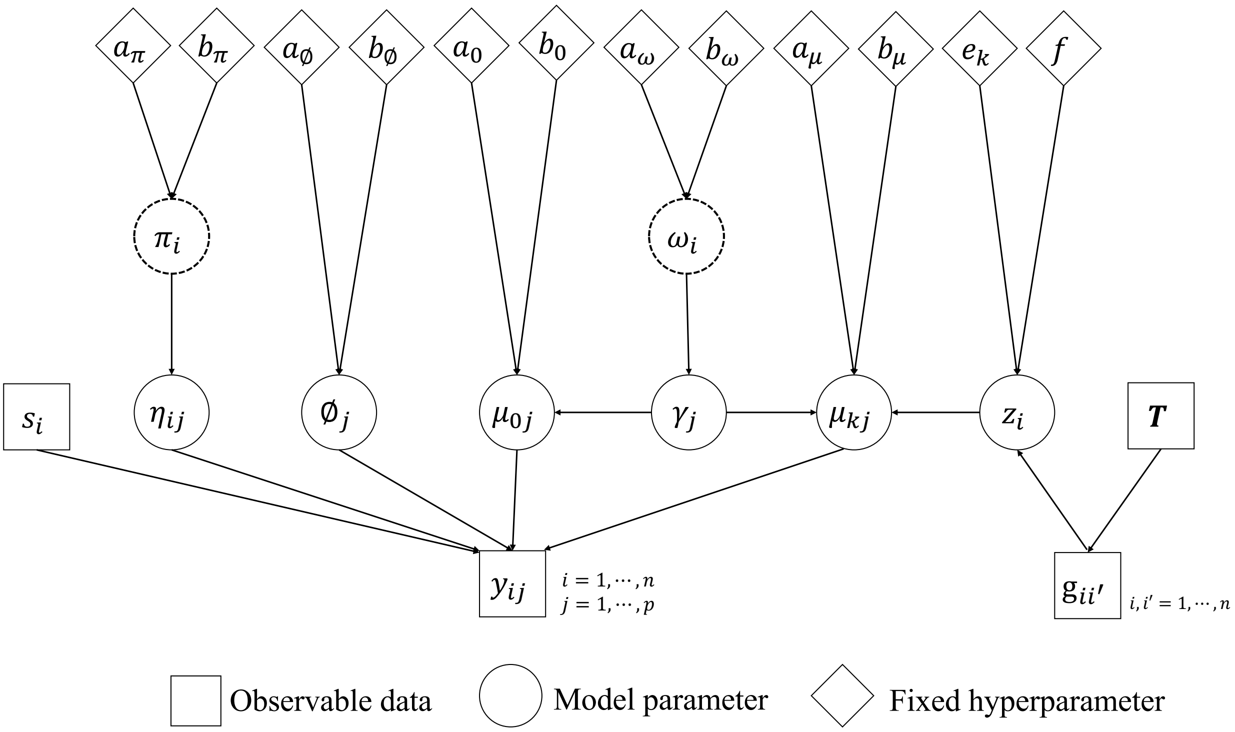


**Figure S1**: Graphical representation of the proposed BayesCafe model. Square, circle, and diamond-shaped nodes refer to observable data, model parameter, and fixed hyperparameter, respectively. Circles with a dashed outline indicate nuisance parameters that are integrated out in BayesCafe. A link between two nodes represents a direct probabilistic dependence.

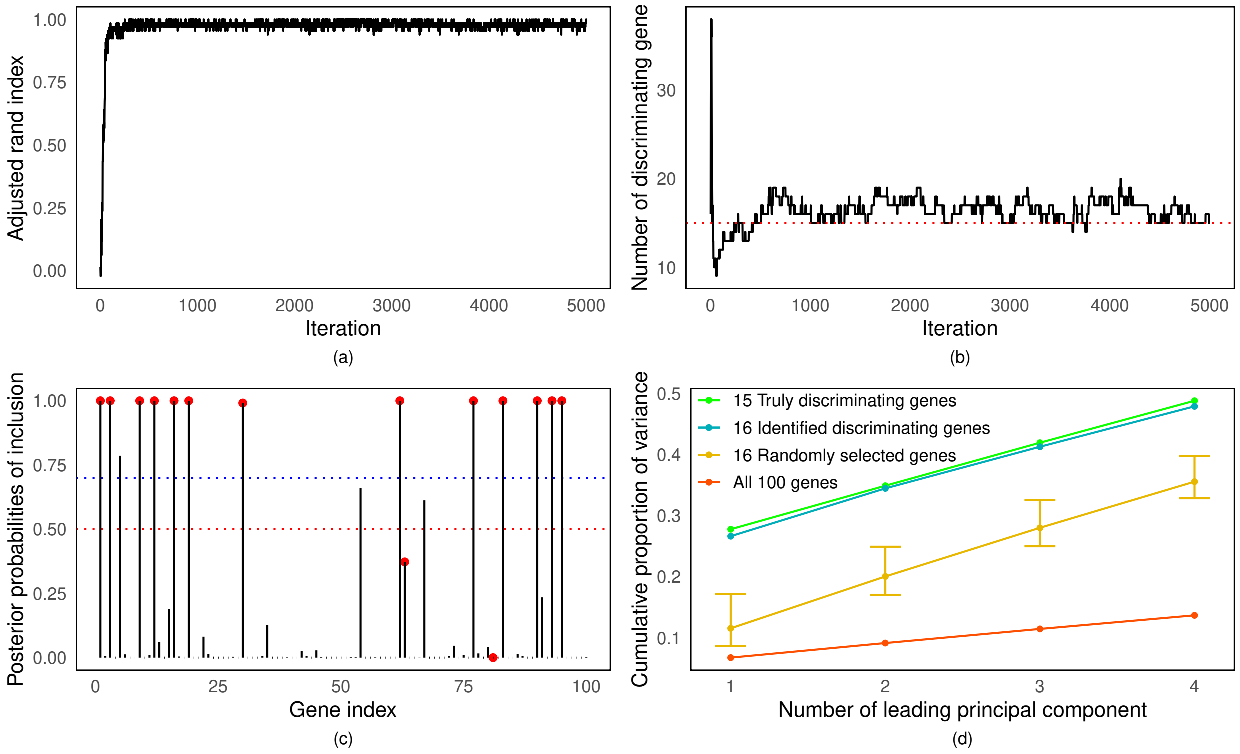


**Figure S2**: The simulation study. (a) The trace plot of the adjusted rand indices (ARI). (b) The trace plot of the number of discriminating genes $p_\gamma$. (c) The marginal posterior probabilities of inclusion of all genes $\Pr(\gamma_j = 1|\cdot)$, with red dots indicating the truly discriminating genes, the horizontal red dotted line indicating a threshold of 0.5, and the horizontal blue dotted line indicating a threshold for a 5% Bayes false discovery rate (BFDR). (d) The cumulative proportion of variance explained by the 15 truly discriminating genes, the 16 discriminating genes identified by BayesCafe, 16 randomly selected genes from all 100 genes (ranges indicated by error bars), and all 100 genes.

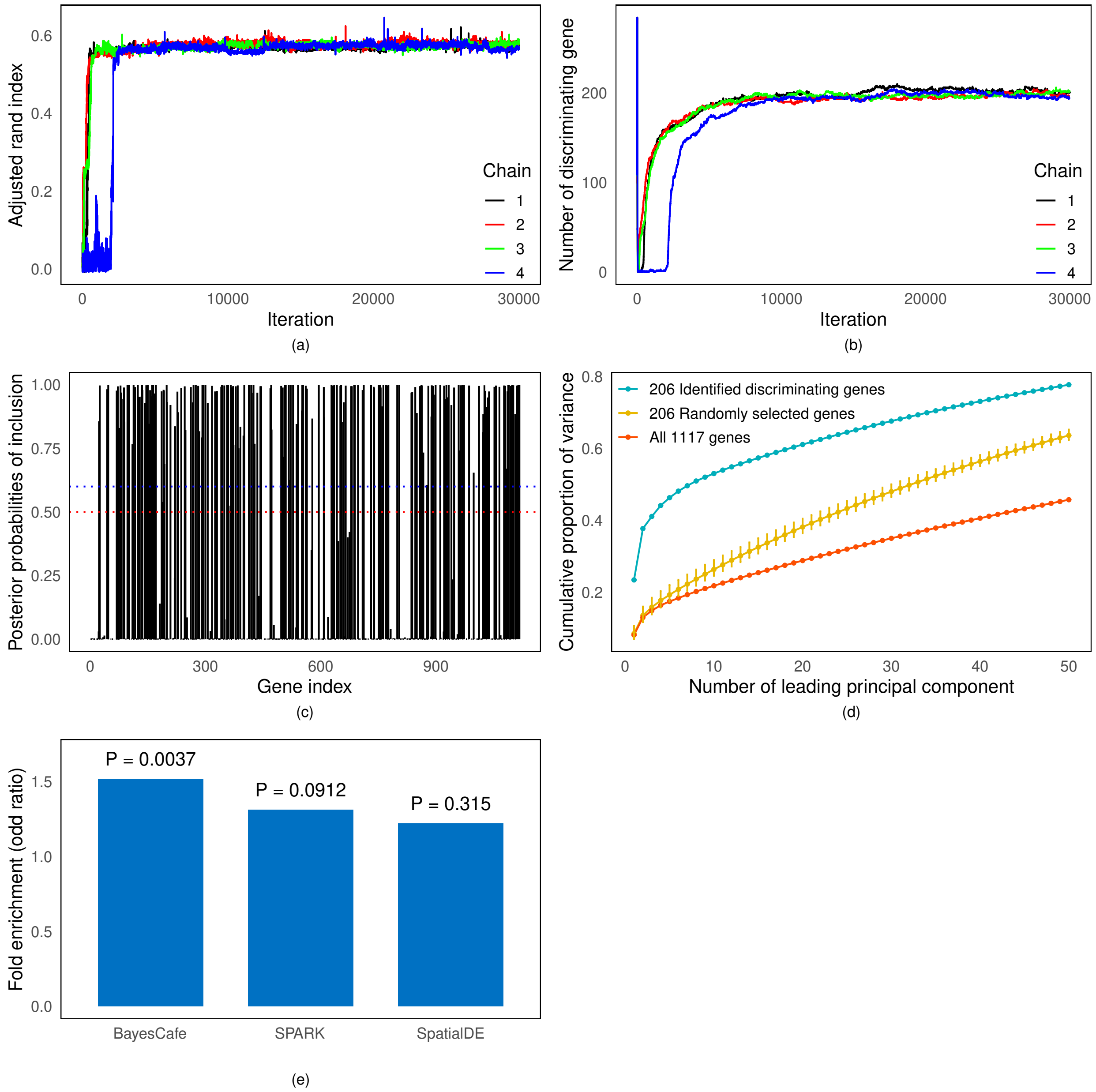


**Figure S3**: The mouse olfactory bulb (MOB) data analysis. (a) The trace plot of the adjusted Rand indices (ARI) of four independent Markov chains. (b) The trace plot of the number of discriminating genes $p_\gamma$ of four independent Markov chains. (c) The marginal posterior probabilities of inclusion of all genes $\Pr(\gamma_j = 1|\cdot)$, with the horizontal red dotted line indicating a threshold of 0.5, and the horizontal blue dotted line indicating a threshold for a 5% Bayes false discovery rate (BFDR). (d) The cumulative proportion of variance explained by the 206 discriminating genes identified by BayesCafe, 206 randomly selected genes from all 1117 genes (ranges indicated by error bars), and all 1117 genes. (e) Bar chart, with the p-values, of fold enrichment values for SV genes detected by BayesCafe, SPARK, and SpatialDE, by querying with the Harmonizome database.

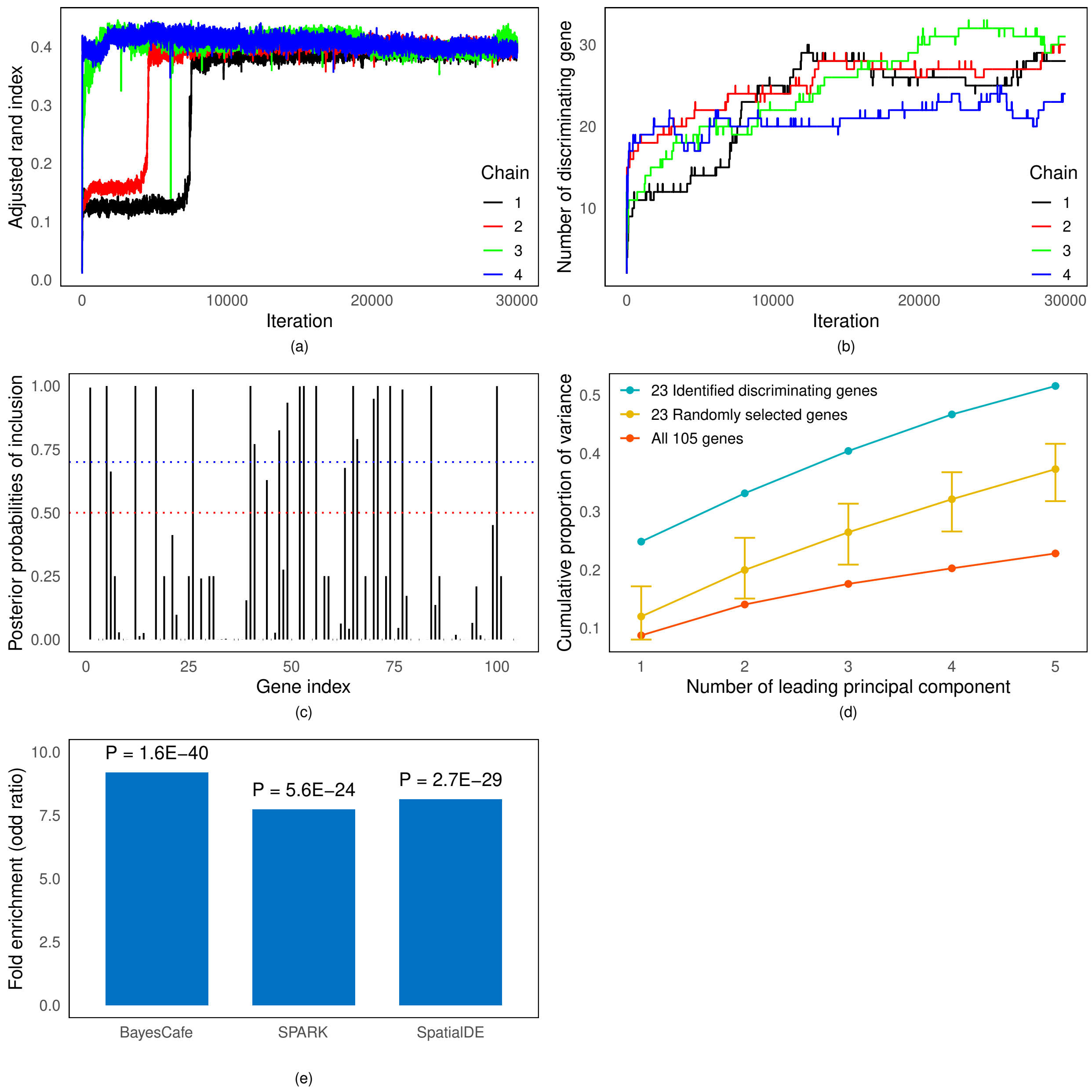


**Figure S4**: The mouse visual cortex data analysis. (a) The trace plot of the adjusted Rand indices (ARI) of four independent Markov chains. (b) The trace plot of the number of discriminating genes $p_\gamma$ of four independent Markov chains. (c) The marginal posterior probabilities of inclusion of all genes $\Pr(\gamma_j = 1|\cdot)$, with the horizontal red dotted line indicating a threshold of 0.5, and the horizontal blue dotted line indicating a threshold for a 5% Bayes false discovery rate (BFDR). (d) The cumulative proportion of variance explained by the 23 discriminating genes identified by BayesCafe, 23 randomly selected genes from all 105 genes (ranges indicated by error bars), and all 105 genes. (e) Bar chart, with the p-values, of fold enrichment values for SV genes detected by BayesCafe, SPARK, and SpatialDE, by querying with the Allen Mouse Brain Atlas.

Table S1: Hierarchical formulation of the proposed BayesCafe model.

**Hierarchical model**:

$$y_{ij}|z_i = k, \eta_{ij}, \phi_j, \gamma_j, \mu_{kj}, \mu_{0j} \sim \begin{cases} 0 & \text{if } \eta_{ij} = 1 \\ \text{NB}(s_i\mu_{kj}, \phi_j) & \text{if } \eta_{ij} = 0 \text{ and } \gamma_j = 1 \\ \text{NB}(s_i\mu_{0j}, \phi_j) & \text{if } \eta_{ij} = 0 \text{ and } \gamma_j = 0 \end{cases}$$

**Cluster Markov random field prior**:

$$\pi(z_i = k|\boldsymbol{z}_{-i}) \propto \exp\left(e_k + f\sum_{i'=1}^{n} g_{ii'}I(z_{i'} = k)\right)$$

**False zero indictor prior**:

$$\eta_{ij}|\pi_i \sim \text{Bern}(\pi_i)$$
$$\pi_i \sim \text{Beta}(a_\pi, b_\pi)$$

**Negative binomial dispersion prior**:

$$\phi_j \sim \text{Ga}(a_\phi, b_\phi)$$

**Feature selection prior**:

$$\gamma_j|\omega \sim \text{Bern}(\omega)$$
$$\omega \sim \text{Beta}(a_\omega, b_\omega)$$

**Normalized expression level prior**:

$$\mu_{kj} \sim \text{Ga}(a_\mu, b_\mu)$$
$$\mu_{0j} \sim \text{Ga}(a_0, b_0)$$

**Fixed hyperparameters**:

$$e_k, f, a_\pi, b_\pi, a_\phi, b_\phi, a_\omega, b_\omega, a_\mu, b_\mu, a_0, b_0$$
